# Functional validation and characterization of *Slc39a8* genotype effects on ethanol consumption and related phenotypes

**DOI:** 10.64898/2026.09.24.754187

**Authors:** André Lucas Silva-Borges, Donna-Jael G. Paredes, Rebeka Sultana, Cecilia G. Pareja, Amanda M. Barkley-Levenson

## Abstract

Alcohol use disorders are heritable, and large-scale genome-wide association studies have recently identified novel genetic loci that are correlated with alcohol consumption and problematic alcohol use. One such gene is *Slc39a8*, which encodes the ion transporter ZIP8, responsible for cellular import of zinc and other metals. *Slc39a8* is highly pleiotropic, and a common missense variant has been associated with decreased problematic alcohol use. Here, we used *Slc39a8* heterozygous (HET) knockout mice and their wild type (WT) littermates to characterize the effects of reduced gene function on two models of voluntary alcohol consumption (drinking in the dark and intermittent access two-bottle choice), and anxiety- and depression-like behavior. We found that HET mice consumed less alcohol than WT in the intermittent access two-bottle choice procedure. No evidence was found for genotypic differences in baseline locomotor activity, anxiety-like, or depression-like behavior in the open field and forced swim tests. *Slc39a8* mRNA expression and alcohol metabolism experiments confirmed decreased brain expression of *Slc39a8* in HET mice and unchanged alcohol metabolism. Basal and post-alcohol measurement of zinc levels showed that HET mice after ethanol exposure had lower zinc concentrations in the liver and higher concentrations in blood than WT animals. Analysis of the spatial distribution of multiple trace metals in the brains of naïve *Slc39a8* HET and WT mice using Laser Ablation Inductively Coupled Plasma Mass Spectrometry (LA-ICP-MS) showed region-specific genotypic differences in levels of zinc and iron. These results provide evidence for a direct role of *Slc39a8* in ethanol consummatory behavior that does not appear to be due to metabolism, affective state, or locomotor activity. Moreover, we confirm that this animal model shows tissue- and brain region-specific alterations in zinc levels, highlighting potential mechanisms for future study.

## Introduction

Alcohol use disorder (AUD) is a major public health concern, with 27.9 million people estimated to meet the diagnostic criteria for an AUD (Substance Abuse and Mental Health Services Administration [SAMHSA], 2025). Additionally, excessive alcohol consumption generates high economic costs, and alcohol is the third leading cause of preventable death in the United States. AUD arises from a combination of genetic and environmental factors, with heritability estimated at approximately 50% (e.g. Verhulst et al., 2015). However, the specific genetic factors that may underlie increased risk for or protection from developing an AUD remain largely unknown. Genome-wide association studies (GWAS) have proven to be an important tool for identifying novel variants and genetic loci implicated in complex traits such as AUD and related behaviors. Recent GWAS of problematic alcohol use and alcohol consumption have identified variants in numerous genes that are correlated with these phenotypes. While this is a crucial first step in uncovering the genetics of AUD, follow-up studies are needed to determine whether these genes play a causal role in alcohol-related traits.

The gene *Slc39a8* has been repeatedly identified in GWAS as associated with problematic alcohol use and number of drinks per week (Sanchez-Roige et al., 2019; Brazel et al., 2019; Liu et al., 2019; Evangelou et al., 2019). This gene encodes the ZIP8 protein, which is a metal cation transporter that functions to import zinc (Zn^2+^), manganese (Mn^2+^), iron (Fe^2+^), and other metals across the cell membrane (Liu et al., 2009). *Slc39a8* is highly pleiotropic and has been associated with multiple other traits including other psychiatric disorders such as schizophrenia (Pardiñas, 2018) and major depressive disorder (Yin, 2022), other substance use disorder traits (Deak et al., 2022; Xu et al., 2023), and physiological traits such as hypertension (Ehret, 2011), liver disease (Parisinos 2020), inflammatory bowel disease (Su et al., 2023), and body mass index (Locke et al., 2015). *Slc39a8* function also appears to be crucial during development, as homozygous knock out of this gene is embryonic lethal (Wang 2011). Such variety of associations highlights the ubiquitous role of *Slc39a8* function across multiple body systems.

A common missense variant in *Slc39a8*, rs13107325 (A391T), results in reduced protein function (Tseng et al., 2021) and has been associated in GWAS with a decrease in alcohol consumption and problematic use (Sanchez-Roige et al., 2019; Brazel et al., 2019; Liu et al., 2019; Evangelou et al., 2019). Importantly, this same variant has also been shown to alter homeostatic maintenance of trace metals (Haller et al., 2018), which represents one possible mechanism by which *Slc39a8* variants may alter physiological and psychiatric traits. For example, Zn^2+^ has been suggested to play a role in both the pathophysiology (Swardfager et al., 2013) and treatment (Szewczyk et al., 2002) of depression. In addition, Zn^2+^ supplementation has been shown to increase alcohol dehydrogenase activity in the liver (Zhou et al., 2005), and Zn^2+^ chelation was shown to prevent alcohol-induced increases of dopamine in the nucleus accumbens (Morud et al., 2015).

However, very little is known about how *Slc39a8* directly affects alcohol-related phenotypes. In *C. elegans*, loss-of-function mutations resulted in a loss in the sedative effects of alcohol on locomotion (Thompson et al., 2020). In contrast, overexpression of this gene in *Drosophila melanogaster* resulted in an increase in alcohol-induced locomotion, whereas a knock down resulted in increased alcohol-induced sedation. In mouse studies, investigation of voluntary drinking found no effect of a heterozygous *Slc39a8* hypomorph mutation on alcohol intake in a continuous access two-bottle choice procedure (Banna et al., 2022). These findings confirm the complexity in the action of this gene in alcohol-related traits, and highlight the need for further investigation in vertebrate models.

Hence, the goal of this study was to characterize the alcohol (ethanol) drinking phenotype of a previously untested line of mice with a heterozygous knockout of *Slc39a8* (HET) compared to their wild type (WT) littermates. We used the drinking in the dark (DID) paradigm to investigate binge-like consumption of ethanol, as well as non-caloric (saccharin) and caloric (sucrose) sweet tastant solutions, followed by a mult-week intermittent access two-bottle choice ethanol drinking (IA-2BC) procedure. We also tested two-bottle choice ethanol DID using the continuous recording BioDAQ system to check for possible differences in drinking microstructure. Given the direction of effect from GWAS where the known *Slc39a8* reduced function variant has been repeatedly associated with decreases in ethanol consumption (Sanchez-Roige et al., 2019; Brazel et al., 2019; Liu et al., 2019; Evangelou et al., 2019), we hypothesized that *Slc39a8* HET mice would have lower ethanol intake in both DID and IA-2BC procedures. We also tested ethanol elimination following ethanol injection to assess potential changes in metabolism. As *Slc39a8* has been associated with other psychiatric traits, and AUDs are frequently comorbid with other psychiatric disorders, we investigated anxiety-like and depression-like behaviors in the open field and forced-swim tests, respectively, in naïve animals to assess potential underlying affective states and locomotor differences that could influence drinking behavior. To begin to investigate essential metals homeostasis as a potential mechanism of *Slc39a8* mutation behavioral effects, we also measured serum, whole brain, and liver levels of Zn^2+^ in naïve and post-drinking animals, and spatial distribution of brain Zn^2+^, Mn^2+^, and Fe^2+^ in naïve mice.

## Materials and methods

### Animals and husbandry

223 male and female *Slc39a8* HET and WT mice were used in these studies. Mice were obtained from an in-house breeding colony at the University of New Mexico Health Sciences Center that was originally established from cryorecovered breeders of strain B6;129S5-Slc39a8^tm1Lex^/Mmuc (stock # 032588-UCD; Tang et al. 2010) purchased from the Mutant Mouse Resource and Research Center (Davis, CA) and maintained with HET x HET breeder pairs. Only HET and WT animals were obtained and used in these experiments, as homozygous knockout of this gene is believed to be embryonic lethal (Wang et al., 2011). Mice were genotyped via ear punch at the time of weaning (Transnetyx, Memphis, TN) and all mice were between 75 and 125 days of age and experimentally naïve at the start of testing. For the drinking experiments (Experiment 1 & 2), animals were singly-housed on a reverse light-dark cycle with lights off at 8:00 am. For all other experiments, mice were group housed 2-6 per cage on a forward light-dark cycle with lights on at 6:00 am. All animals were given *ad libitum* access to food (5L0D LabDiet) and water. All procedures were approved by the University of New Mexico Health Sciences Center Institutional Animal Care and Use Committee and were conducted in accordance with the NIH Guidelines for Care and Use of Laboratory Animals.

### Drugs

Ethanol (200 proof, Decon Laboratories, King of Prussia, PA) for all drinking experiments was diluted in tap water at the concentration of 20% (v/v). For the ethanol metabolism experiment, ethanol solution was prepared with 0.9% saline (20% v/v) and intraperitoneally administered at a dose of 2 g/kg. Saccharin (Sigma-Aldrich, Saint Louis, MO) was diluted in tap water at the 3.2mM concentration. Sucrose (Gibbstown, NJ) was diluted at the concentration of 10% in tap water.

### Drinking in the Dark (DID)

During the DID procedure, mice were weighed approximately 30 min prior to the start of the drinking sessions. Three hours into the dark cycle, water bottles were removed and replaced by 10 ml tubes fitted with a ball bearing metal sipper and containing the solution to be tested. Fluid levels were recorded, and tubes were left in place for 2 hours, after which fluid levels were recorded again. Tubes were then removed and replaced by water bottles. On Day 4, the drinking tubes were left in place for 4 hours, and fluid levels were recorded after 2 and 4 hours. Average fluid level readings obtained from two empty control cages per rack were used to adjust the recorded intake levels for spillage and evaporation.

### Intermittent Access Two-Bottle Choice Drinking (IA-2BC)

In this paradigm, animals received 24-hour access to two drinking tubes 3 times a week: one containing 20% (v/v) ethanol in tap water and the other containing tap water. This access occurred on alternating days (i.e., every other day) for a total of 3 days a week for three consecutive weeks. On each ethanol access day, mice were weighed, and their standard water bottles were removed and replaced with the two experimental tubes. The position of the ethanol and water tubes was alternated on each access day to avoid the development of a side preference. On intervening non-drinking days, a single water bottle was reintroduced and placed in the central position of the home cage. Average fluid level readings obtained from two empty control cages per rack were used to adjust the recorded intake levels for spillage and evaporation.

### Continuous Fluid Consumption Recording

In Experiment 2, the BioDAQ Episodic Intake Monitoring System was used to continuously record fluid consumption, as has been previously reported (Barkley-Levenson and Crabbe, 2012). Mice were housed in specialized polycarbonate shoebox cages containing seven cutouts for the attachment of recessed liquid hoppers that are connected via ethernet cables to the BioDAQ recording system. Access to each hopper can be controlled manually by opening or closing a gate that sits flush with the side of the cage. During the acclimation period and when animals were not being tested, a liquid hopper containing water was attached to the center position on the side of the cage and access gates to the other hopper positions were closed. During testing, both water and ethanol hoppers were attached to the left and right positions on the side of the cage and the center hopper position was closed. Fluid intake was measured with force sensors that record the weight of the bottles once every second and register intake with a sensitivity of 0.01g when the animal applies a 0.08g or greater force to the hopper.

### Open Field Apparatus

The open field apparatus consisted of an arena made of acrylic transparent walls and white floor (27.3 x 27.3 x 20cm; Med Associates, St. Albans, VT). This arena was placed inside of a light- and sound-attenuating cubicle with inside illumination. To detect horizontal and vertical activity, the chamber was equipped with infrared light sensors separated by 1.5 cm and at 1.5 and 6 cm above the floor level, respectively. The center space was defined as 8×8 beams (14.45 cm x 14.45 cm). Locomotor activity (distance traveled in cm) and the time spent in the center of the arena were automatically recorded.

### Experiment 1: Ethanol and sweet tastant DID tests followed by IA-2BC

Before the start of the experiment, mice were single-housed and allowed to acclimate to a reverse light-dark cycle for 2 weeks. This experiment was divided into 4 phases. Phases 1 to 3 consisted of DID sessions of different substances: ethanol (20% v/v in water), saccharin (3.2 mM) and sucrose (10% w/v in water). A 10-day washout period was given between ethanol and saccharin DID, and at least three days were given between the end of the saccharin DID and the start of the sucrose DID. Phase 4 (IA-2BC) started at least three days after the last day of sucrose DID. During this phase, mice received 3 weeks of IA-2BC ethanol drinking as described above. After the final day of drinking, animals were sacrificed via carbon monoxide asphyxiation and brain, blood, heart, and livers were removed from a subset of animals in the experiment.

### Experiment 2: Two-bottle choice DID and drinking bout microstructure

Animals in this experiment were allowed to acclimate to a reverse light-dark cycle and to single-housing conditions for 2 weeks. On the first day of the experiment, animals were weighed prior to the start of the drinking session at approximately 3 hours into the dark cycle. The water and ethanol bottles were placed on their respective hoppers, hopper gates were opened, and recording was initiated for each session. Except for the use of an automated system for measurements and the addition of a bottle with water during the experiment, the remainder of this experiment followed the DID protocol described above. Only one week of DID was conducted, with 3 days of 2h access to ethanol and water, and a final extended day of access where water and ethanol bottles were left in place and intake was continuously recorded for 24 hr. Intake was measured during the first 4 hr for comparison with standard homecage DID methods, and the total 24 hr intake was also measured to determine potential genotypic differences that emerged with prolonged ethanol access (similar to an individual drinking session in the IA-2BC procedure).

### Experiment 3: Open field test (OFT) and forced swim test (FST)

Animals were moved to the experiment room and allowed one hour for acclimation. Mice were then placed individually into the center of the OFT arena and given 30 minutes to explore it freely. Locomotor activity (distance traveled in cm) and the time spent in the center of the arena were automatically recorded. Between animals, the chambers were cleaned with 10% isopropyl ethanol to eliminate odor cues. Two days after the OFT, the animals were moved to a new experiment room and were allowed 30 minutes for acclimation. Following this period, mice were placed into 14×23cm glass cylinders filled halfway with water at the range of 23-25° Celsius, where they were given a 6-minute forced swim test. Mice were video recorded during this period from the side of the glass. Water was replaced between each animal. Once the test was finished, the final 4 minutes of the video were analyzed and scored for time spent immobile by an experimenter blinded to sex and genotype.

### Experiment 4: Ethanol Metabolism

In order to assess the rates of ethanol clearance, mice were injected with 2 g/kg of ethanol intraperitoneally. Retro orbital blood samples were then taken at different time points depending on the group assigned: Group 1) 15, 60, 180 minutes or Group 2) 30, 120, 240 minutes post-injection. Blood samples (20µL) were taken alternating the eyes (left and right), with the final sample taken from the same eye as the first sample. Blood samples were transferred to centrifuge tubes and left in ice until processing after all samples had been collected. To process these samples, the tubes were centrifuged at 3000 rpm for 10 minutes and the supernatant was then collected and transferred to a new tube to be used for BEC analysis in an Analox Alcohol Analyzer (Analox Instruments, Lunenburg, MA).

### Experiment 5: Phenotypic characterization of trace metal levels and Slc39a8 gene expression in HET and WT mice

Because of ZIP8’s role as a putative Zn^2+^ transporter, we chose to characterize potential changes at baseline and following ethanol intake in blood and tissue (brain and liver) levels in naïve mice and a subset of mice following the end of the IA-2BC drinking in Experiment 1. Zn^2+^ levels in blood serum, whole brain, and liver were measured using a colorimetric assay kit (Abcam, ab102507) following manufacturer’s protocols. Briefly, the tissues were homogenized and lysed in a PBS + 0.1% TritonX-100 (Sigma-Aldrich, Saint Louis, MO) buffer and deproteinized with 7% TCA solution buffer. After deproteinization, the supernatants were separated and mixed with the Zinc Reaction Buffer in a 96-well plate. After a 10-min incubation period, the plates were placed in a microplate reader and measured at OD_560_. The final concentration values were calculated from the zinc standard curve after subtracting background values. Laser Ablation Inductively Coupled Plasma Mass Spectrometry (LA-ICP-MS) was used for semi-quantitative analysis of the spatial distribution of Zn^2+^, Mn^2+^, and Fe^2+^ in brains from 4 HET and 2 WT naïve mice. After collection, brain samples were sagittally divided and right hemispheres were sectioned in a cryostat at 50 µm thickness starting at the midline and slices at approximately 0.675mm lateral coordinates were mounted on gelatin-coated glass slides. These slides were then analyzed by the University of New Mexico Integrative Molecular Analysis Core using an imageBIO266 laser ablation system (Elemental Scientific Lasers, Bozeman, MT, USA) that includes a 266nm wavelength, 7J/cm2 energy fluence, 10Hz repetition rate, 50μm spot size, 500 μm/s scan speed, and a helium (He) carrier gas flow rate of 0.8 L/min. This equipment is coupled with a Dual Concentric Injector (DCI) connected to an Agilent 7900 ICP-MS equipped with an RF power of 1550W, a nebulizer gas (Ar) flow rate of 0.44 L/min, and a dwell time of 10ms.

In a separate set of animals, *Slc39a8* mRNA levels were measured in whole brain samples from naïve mice to confirm reduced gene expression in the HET mice. Briefly, whole brains were homogenized and approximately 30mg were separated and the RNA content was isolated using a QIAshredder and RNeasy Kit following manufacturer instructions (Qiagen, Germantown, MD). Then, the isolated RNA was used as template to prepare cDNA with a High-Capacity cDNA Reverse Transcription Kit (ThermoFisher Scientific, Waltham, MA).

For the qPCR preparation, *β-actin* and *Slc39a8* probes (ThermoFisher Scientific, Waltham, MA) were used and the reaction was setup using Universal Master Mix (ThermoFisher Scientific, Waltham, MA). The final product was ran using triplicates and analyzed in a BioRad CFX384 Touch Real-Time PCR Detection System (BioRad, Hercules, CA). Final gene expression levels were calculated with the CT method using *β-actin* as the endogenous control.

### Statistical analyses

Statistical analyses were carried out using IBM SPSS Statistics software (version 30.0.0.0; IBM Corporation, Armonk, NY) and graphed with GraphPad Prism software (GraphPad Prism, v. 10.5.0, La Jolla, CA). Unless otherwise noted, sex and genotype were included as factors in all analyses in this study. Because of occasional missing data during drinking periods due, linear mixed models were used to analyze DID and BioDAQ two-hour intake, and IA-2BC drinking data, whereas total values and averages were analyzed via two-way analysis of variance (ANOVA). Body weight data from animals in Experiment 1 were analyzed using repeated measures ANOVA. Two-way ANOVAs were conducted to analyze data on Experiments 3 and 5. For the brain LA-ICP-MS metal levels in Experiment 5, CPS was adjusted dividing the region values by the region of interest (ROI) size in each sample, and the adjusted values were used for statistical analysis. Due to interleaved sampling design on Experiment 4, data was analyzed via linear mixed models. Welch’s t test was used to analyze RT-qPCR data. Data that were ± 2.5 standard deviations from the group mean were considered statistical outliers and removed. Data are shown collapsed on sex when there was no significant interaction between sex and other factors. α=.05 unless otherwise stated. P values less than .08 are discussed as potential statistical trends.

## Results

### Experiment 1: Ethanol and sweet tastant DID tests followed by IA-2BC

Figure 1 shows average g/kg intake of ethanol, and ml/kg of saccharin and sucrose on each day of the different DID cycles (1B, C, and D) and IA-2BC drinking (1E). Water consumption (1F) and preference ratio (1G) during the IA-2BC procedure are also shown. In panels H-J, we show average daily ethanol and water consumption, and preference ratio, respectively. For the average daily preference ratio data, two outliers were removed (1 Female WT, 1 Female HET). No significant differences were found between the genotypes for ethanol (*F*(1,44)= 0.439, *p=*0.511), saccharin (*F*(1,45)=0.011, *p=*0.916) and sucrose (*F*(1,41)=1.011, *p=*0.321) DID when analyzed via repeated measures ANOVA. For the ethanol IA-2BC cycles (Fig. 1E), linear mixed model analysis found a significant effect of genotype (*F*(1, 65.971)= 10.073, *p=*0.002) for ethanol consumption across sessions, with the HET mice exhibiting lower intakes than their WT littermates. There was also a significant effect of sex (*F*(1,65.971)= 65.974, *p<*0.001), but since there was not a significant sex x genotype interaction (*F*(1,37)= 0.100, *p*<0.753), the data are shown collapsed by sex. We found that there was not an effect of genotype on water consumption (Fig. 1F) or preference ratio (Fig. 1G). Daily averages for water consumption and preference ratio also showed no difference between the genotypes (Fig. 1 panels I and J, respectively). When the average daily ethanol consumption was analyzed, however, a two-way ANOVA showed a significant effect of genotype (*F*(1,48)= 6.979, *p=* 0.011) (Fig. 1H).

**Figure 1:**
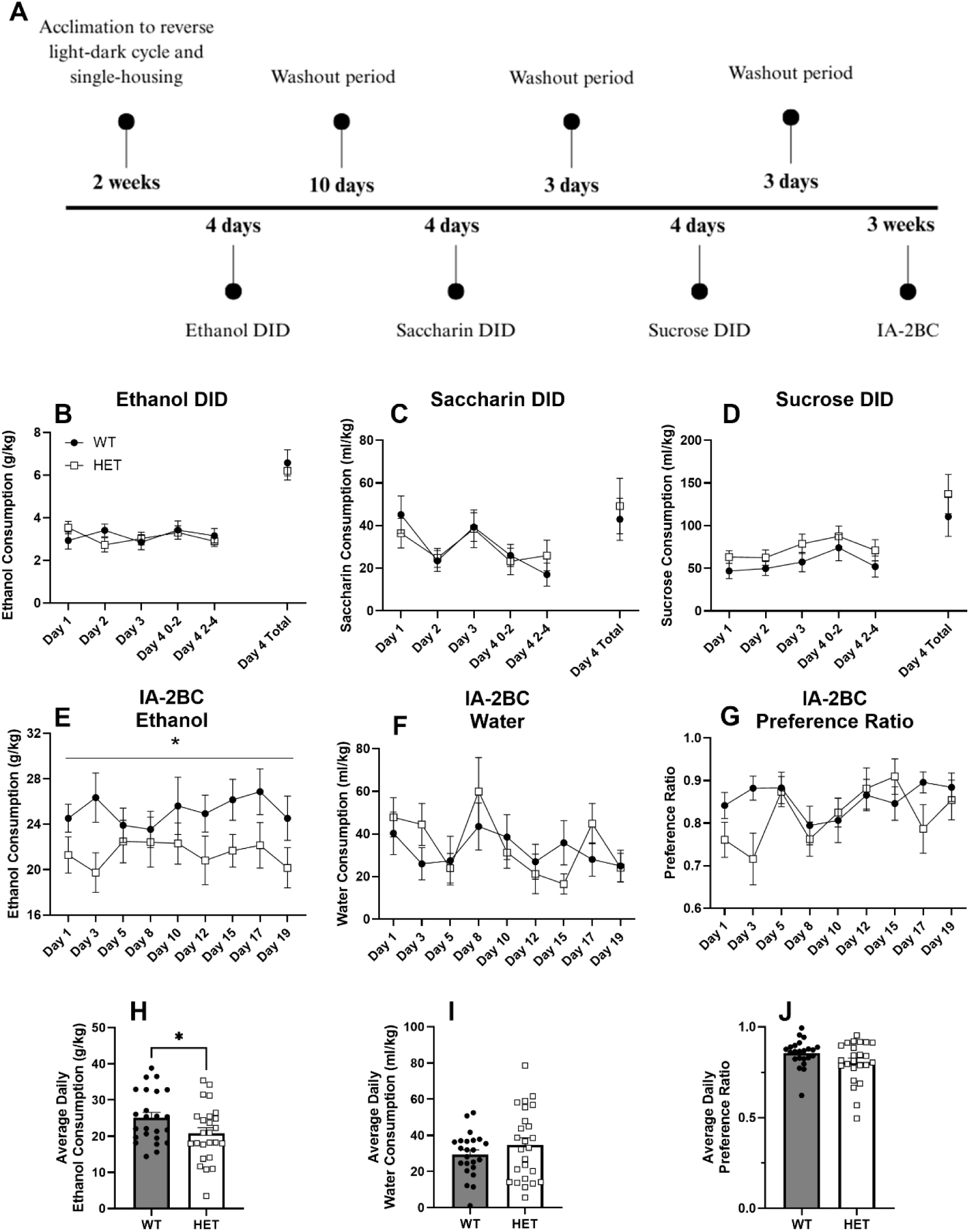
Consummatory behavior of *Slc39a8* HET mice and their WT littermates. Panels show experiment timeline (A), ethanol (B), saccharin (C), and sucrose (D) consumption during DID, ethanol (E) and water consumption during IA-2BC (F), preference ratio of ethanol during IA-2BC (G), and average daily consumption of ethanol (H) and water (I) and average preference ratio (J) during IA-2BC. Data are shown as mean ± SEM. There were no statistically significant differences between the genotypes for consumption during DID trials. There was a significant difference between the genotypes for IA-2BC ethanol consumption. * Indicates *p<0.05.* N= 11-13/sex/genotype

In Experiment 1, the animals were weighed on each experimental day, and we chose to analyze body weight data for a potential genotypic difference due to the previous association of *Slc39a8* variants with body mass index in human GWAS (Locke et al., 2015). Weights on each experimental day are shown in Figure 2. Repeated measures ANOVA showed a significant effect of genotype (*F*(1,37)= 8.294, *p*= 0.007), a main effect of sex (*F*(1,37)= 25.277, *p*<0.001, and no sex x genotype interaction (*F*(1,37)= 0.096, *p*= 0.759), with the HET animals showing higher body weight across the experiment than the WT mice.

**Figure 2:**
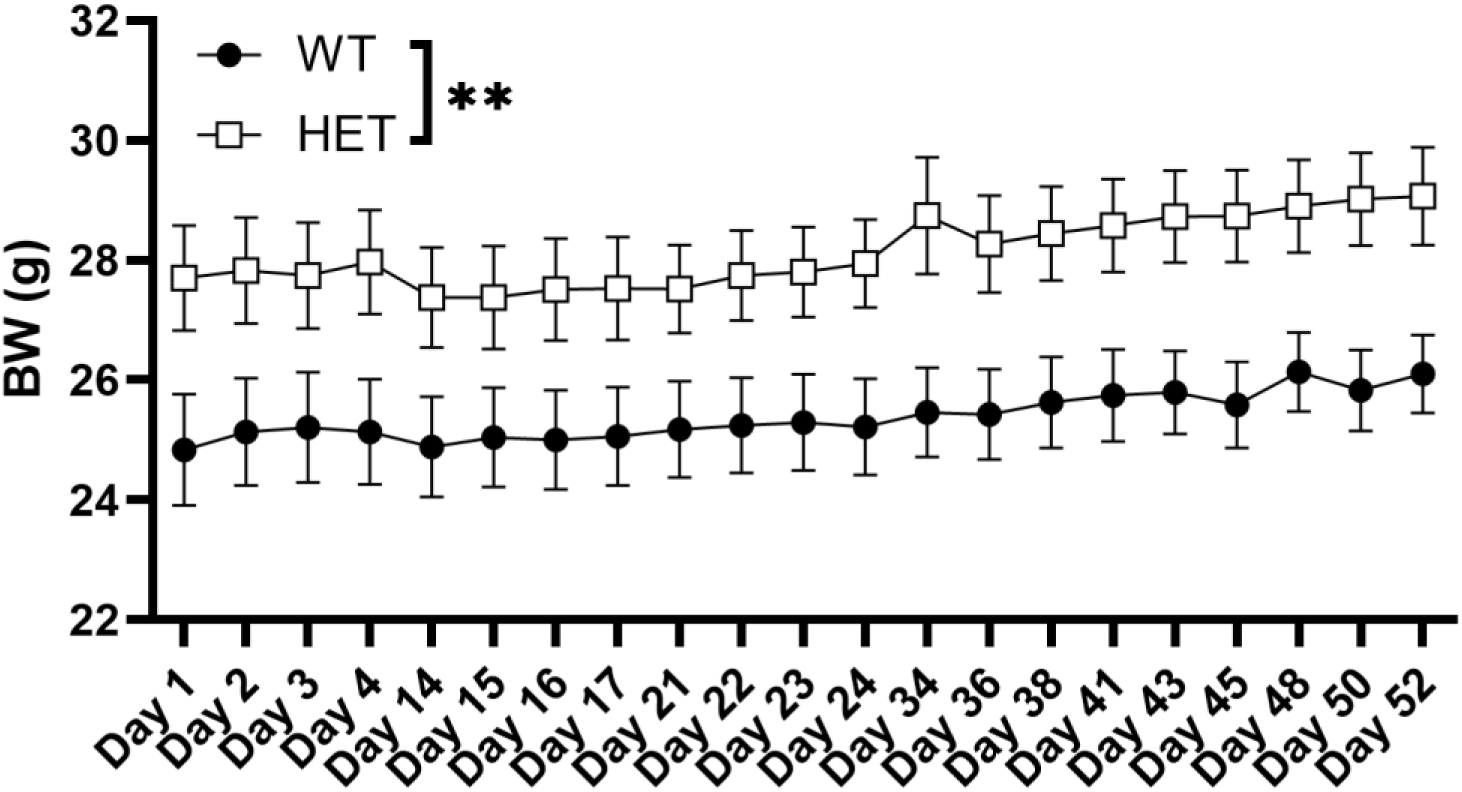
Body weight progression of *Slc39a8* HET mice and their WT littermates during DID and IA-2BC drinking in Experiment 1. Data are shown as mean ± SEM. ** indicates *p<0.01.* N= 11-13/sex/genotype.

### Experiment 2: Two-bottle choice DID and drinking bout microstructure

To assess drinking microstructure and whether the presence of a choice between water and ethanol could explain the lower intake seen in HET mice during IA-2BC, we utilized the continuous recording BioDAQ apparatus to investigate drinking rate across the sessions. Cumulative intake over time can be seen in Supplementary Figure S1. We did not see any significant genotypic differences for ethanol (Figure 3A) or water (Figure 3B) consumption (ethanol: (*F*(1, 9) = 0.5629, *p*= 0.47); water: (*F*(1, 9) = 0.00023, *p*= 0.998)). Due to low sample number per sex (n=2-3/sex/genotype), data were analyzed without sex as a variable. We also did not find significant differences for ethanol preference or drinking microstructure variables across sessions, including number of bouts, intake rate, average bout size, and interbout interval (Supplementary Figure S2).

**Figure 3:**
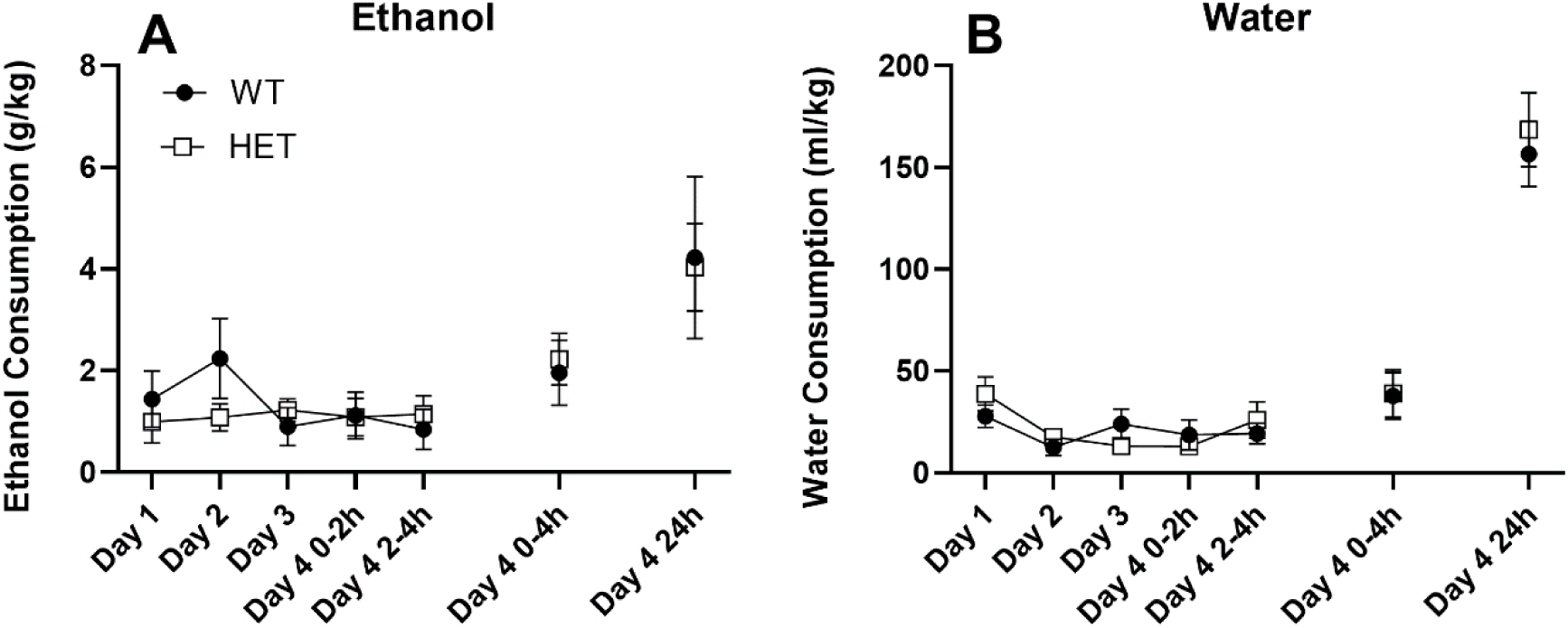
Two-bottle choice DID in the BioDAQ. Panels show ethanol (A) and water (B) consumption during experiment days. Data are shown as mean ± SEM. N= 5-6/genotype.

### Experiment 3: Open field test (OFT) and forced swim test (FST)

For the OFT, we analyzed percentage of the total time spent in center and the total distance traveled over the 30 min test. Figure 4A shows the percentage of time spent in the center. No significant differences between the groups were found, (*F*(1,91)= 0.010, *p*= 0.919). There were also no statistically significant differences between the genotypes for the distance traveled (*F*(1,91)= 0.348, *p=* 0.557), as seen in figure 4B, and for the time spent immobile during the FST (*F*(1,90)= 0.025, *p=* 0.875), showed in 4C. For all variables analyzed, there were also no main effects of sex, and no significant sex x genotype interactions (p>0.05 for all). Together, these findings suggest there are no baseline differences for anxiety-like or depression-like behaviors between the genotypes.

**Figure 4:**
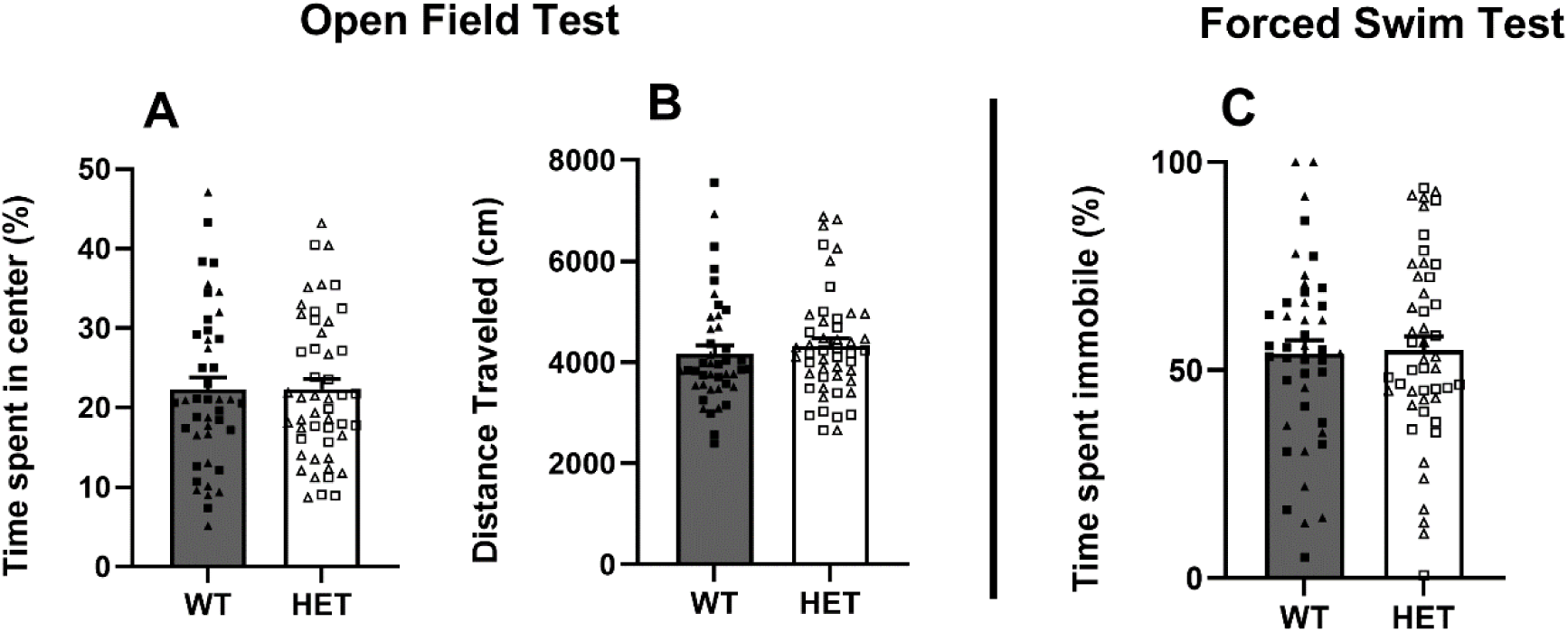
Panels show percent time spent in center (A) and distance traveled (B) during OFT, and time spent immobile (C) during FST. Means ± SEM shown. Males are represented with square and females with triangles, though there were no main effects of sex identified and no significant sex x genotype interactions. N= 19-27 sex/genotype.

### Experiment 4: Ethanol Metabolism

Figure 5 shows an elimination curve depicting average BECs at different timepoints after 2 g/kg ethanol injections. There was no main effect of genotype (*F*(1,27.106)= 0.367, *p=*0.550) or genotype x time interaction (*F*(1,5)= 0.286, *p*= 0.919), no main effect of sex (*F*(1, 27.106)=1.173, *p=*0.288), and no sex x genotype interaction (*F*(1, 27.106)=0.232, *p=*0.634), indicating no difference in ethanol metabolism between the groups.

**Figure 5:**
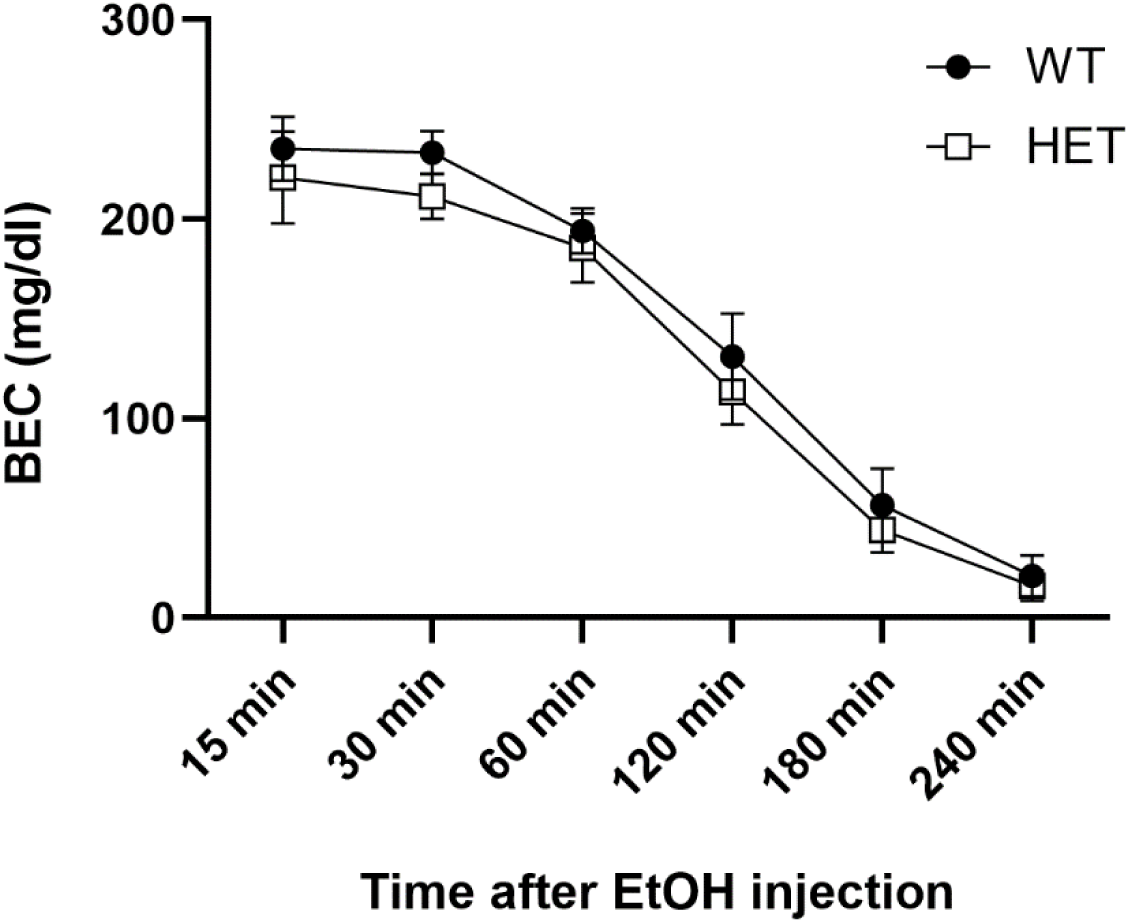
BECs at different timepoints after 2 g/kg (v/v in 0.9% saline solution) injections in *Slc39a8* HET mice and their WT littermates. Data are shown as mean ± SEM. N= 27-33 per genotype per timepoint.

### Experiment 5: Phenotypic characterization of trace metal levels and Slc39a8 gene expression in HET and WT mice

To confirm that our model has diminished brain *Slc39a8* expression, a subset of the naïve animals used in the analysis of brain Zn^2+^ was also used to assess *Slc39a8* mRNA expression (Figure 6). Our results confirmed decreased *Slc39a8* expression in our HET mice relative to WTs (*F*(3,5)= 6.765, *p=* 0.0227)

**Figure 6:**
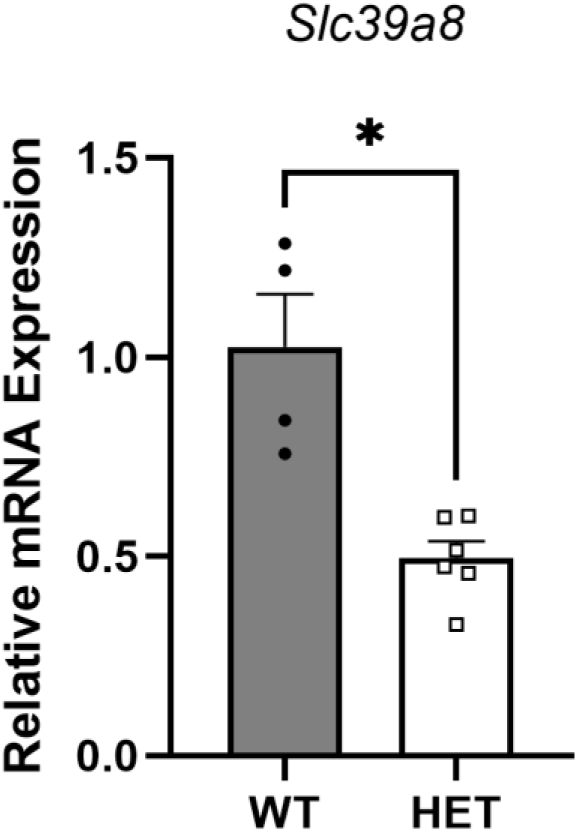
mRNA expression levels of naïve *Slc39a8* HET mice and their WT littermates. Data are shown as mean ± SEM. * Indicates *p<0.05.* N= 4-6 per genotype.

To assess whether there are baseline differences in Zn^2+^ content in brain, liver, and blood of our HET mice in comparison to their WT littermates, samples were taken from adult naïve animals (Fig. 7). There was a significant sex x genotype interaction for the brain Zn^2+^ levels (*F*(1,17)= 4.766, *p=* 0.047), but follow-up ANOVAs for each sex found only a trend towards a genotypic effect in the female mice (*F*(1,7)= 4.561, *p=* 0.077), with the HETs exhibiting higher Zn^2+^ levels than the WTs. There were no other main effects or significant interactions seen for the other tissues. We also analyzed samples taken from two cohorts of mice that underwent the drinking paradigms described in Experiment 1 (Figure 7B). For post-drinking animals, Zn^2+^ levels in our HET mice were found to be lower in the liver (*F*(1,15)= 6.285, *p=* 0.028) and higher in blood (*F*(1,6)= 8.050, *p=* 0.036). No significant difference was found for the brain levels (*F*(1,11)= 1.415, *p=* 0.268). There were no sex x genotype interactions for either liver (*F*(1,15)= 0.143, *p*= 0.712) or brain (*F*(1,11)= 1.226, *p*= 0.300). It is important to note that we were unable to use sex as a factor for the analyses of the blood levels due to the small sample size, and caution is therefore warranted when interpreting these findings.

**Figure 7:**
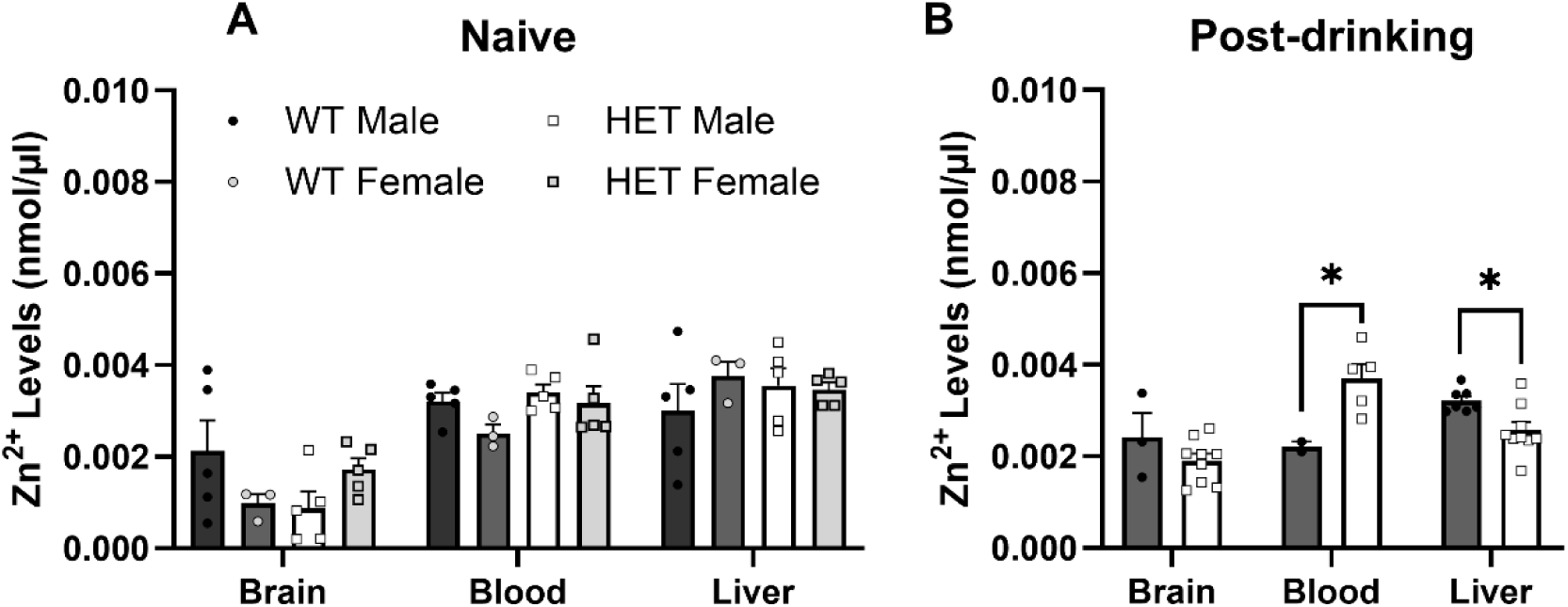
Zinc levels in brain, blood, and liver. Panels show levels from naïve (A) and ethanol-exposed (B) *Slc39a8* HET mice and their WT littermates. Data are shown as mean ± SEM. * Indicates p<0.05. N= 2-10 per genotype depending on the tissue analyze

We hypothesized that brain metal concentrations might show region-specific genotypic differences that were obscured when measuring whole brain levels. To investigate this possibility, we employed LA-ICP-MS to image sagittal brain slices and acquire a semi-quantitative analysis of Zn^2+^, Mn^2+^, and Fe^2+^ in some key regions of interest. The results for Zn^2+^ are shown in Figure 8, and Fe^2+^ and Mn^2+^ concentrations are shown in Supplementary Figures S3 and S4. We found a significant genotype effect on Zn^2+^ content in the ventral striatum (*F*(1,5)= 23.333, *p=* 0.008), and trends toward genotype effects in the prefrontal cortex (*F*(1,5)= 7.522, *p=* 0.052) and hypothalamus (*F*(1,5)= 5.528, *p=* 0.078), with HET levels higher than WT. We found no genotypic difference in Zn^2+^ levels in the hippocampus (*F*(1,5)= 0.506, *p=* 0.516). *Slc39a8* HET brains also showed higher ventral striatal levels of Fe^2+^ (*F*(1,5)= 27.013, *p=* 0.007) compared to WT littermates (Figure S4), but no differences were seen for Mn^2+^ in any region (Figure S3).

**Figure 8:**
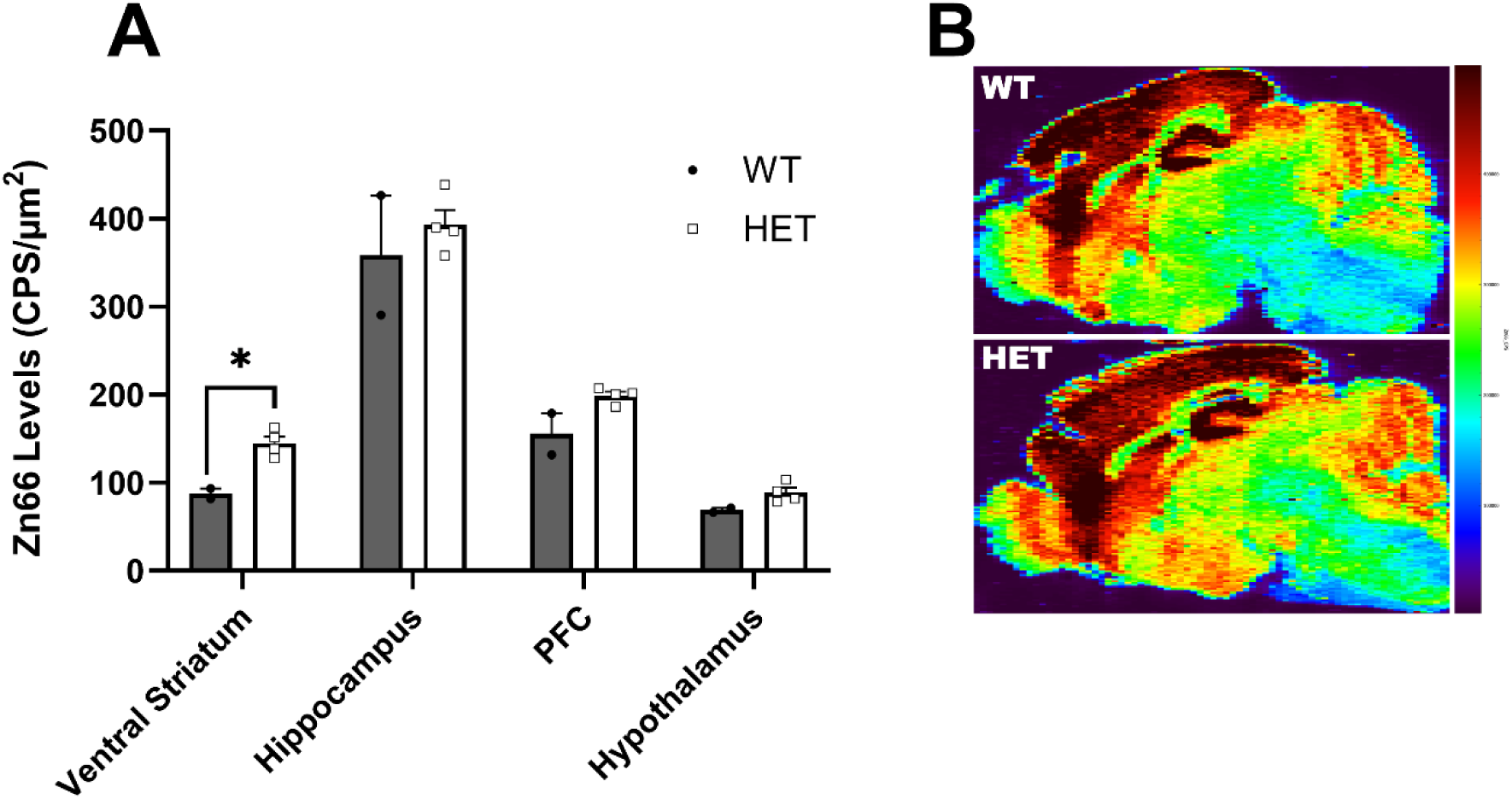
Semiquantitative levels of Zn^2+^ in different brain regions of naïve *Slc39a8* HET mice and their WT littermates (A) and representative elemental distribution map of zinc levels from the LA-ICP-MS analysis (B). Data are shown as mean ± SEM. * Indicates p<0.05. N= 2-4 per genotype.

## Discussion

To follow up on the GWAS finding of an association between *Slc39a8* and problematic ethanol use and drinking, we performed some of the first comprehensive phenotyping of the *Slc39a8* heterozygous constitutive knockout mice for ethanol-relevant traits. We confirmed a role for this gene in intermittent ethanol consumption that does not appear to be explained by metabolic differences or general changes in rewarding fluid consumption. Given the association of this gene with other psychiatric disorders, we also evaluated the *Slc39a8* HET and WT mice for baseline differences in affective behaviors, where no genotypic differences were found. Importantly, reduced brain *Slc39a8* expression in HET mice was confirmed, and baseline and post-ethanol Zn^2+^ concentrations were characterized in both genotypes. In addition, brain regional distribution of metals in naïve mice was also investigated and showed region-specific increases in Zn^2+^ levels in HET mice compared to WT. Taken together, our results provide some of the first evidence supporting a causal relationship between *Slc39a8* expression levels and ethanol consumption.

To further explore the relationship between *Slc39a8* and ethanol intake, we tested *Slc39a8* HET mice on a series of drinking paradigms. When we assessed binge-like ethanol consumption in a 4-day DID model, we saw no significant genotype effects on drinking, with both genotypes consuming similarly large quantities of ethanol. Given the role of sweet taste in voluntary ethanol consumption in mice (Blednov et al., 2008), we also performed saccharin and sucrose DID tests. We found no evidence of genotype effects on saccharin or sucrose binge-like consumption, suggesting that changes in caloric or non-caloric sweet solution consumption do not underlie the differences in ethanol intake in these animals.

Interestingly, our HET mice exhibited lower ethanol consumption in our 3-week IA-2BC paradigm. This is consistent with GWAS evidence linking *SLC39A8* variants that result in reduced protein function to lower alcohol consumption and problematic alcohol use (Sanchez-Roige et al., 2019; Brazel et al., 2019; Liu et al., 2019; Evangelou et al., 2019; Tseng et al., 2021). These findings differ from what was found in a previous study investigating ethanol intake in a different model of *Slc39a8* deficiency (Banna et al., 2022), which did not see any effects of genotype on continuous access two-bottle choice ethanol drinking. In addition to using a different line of *Slc39a8-*deficient mice in our study, there were also several procedural differences that could potentially account for the difference in genotype effect. First, our study used an intermittent access rather than a continuous access drinking procedure. Repetitive cycles of ethanol access and deprivation in the intermittent access model have been show to produce sustained higher levels of ethanol consumption and escalation of preference compared to continuous access models (Hwa et al.,2011), which could potentially reveal genotypic differences not seen at the lower intake levels reported in the previous study. Additionally, the mice in our experiment had prior binge-like ethanol experience, which has been shown to impact subsequent ethanol intake (Cox et al., 2013) and could also contribute to the difference in genotype effects across studies.

In contrast to the results of the IA-2BC drinking results, we did not observe any difference between HET and WT mice in binge-like drinking in the DID test. One possibility for this difference could be the lack of choice in our DID model. To test this possibility, we also examined intake during a two-bottle choice 4-day DID procedure with 20% ethanol vs water in specialized BioDAQ cages that continuously record intake. We also provided extended access on the final day of DID to assess when the genotypes might begin to diverge in intake during a 24-hr access period. We found no genotypic differences in two-bottle choice DID, suggesting that it is not simply the presence of a choice between water and ethanol that leads to reduced ethanol intake in the HET mice. We also did not find any intake differences during the extended access session. It should be noted that mice in Experiment 1 had 24 days between their last DID session and the first 24 hr IA-2BC session, whereas the extended drinking session immediately followed DID in Experiment 2. Mice in Experiment 1 also experienced two other drinking solutions prior to IA-2BC, and because of these procedural differences we are unable to directly compare the findings between the two experiments. Additionally, ethanol intake was reduced in the BioDAQ cages compared to standard homecages, as has been previously reported (Barkley-Levenson and Crabbe, 2012), and this reduced intake could potentially mask genotypic differences in consumption.

Therefore, further studies will be needed to identify which specific factors may contribute to the differences in genotype effect seen between DID and IA-2BC consumption.

*Slc39a8* variants have been linked with other psychiatric disorders including schizophrenia and major depression disorder (Pardiñas et al., 2018; Yin et al., 2022). Additionally, other mental health disorders such as anxiety and mood disorders are frequently comorbid with AUD (Ummels et al., 2022; Grant et al., 2004). Thus, we employed the open field and forced swim tests to investigate affective and locomotor behaviors in naïve animals, and to determine whether there were phenotypic differences that could potentially explain the reduced ethanol intake by HET mice. Our results showed that naive HET mice and their WT littermates have similar locomotor activity and time spent in the center during OFT, as well as no differences in time spend immobile during the FST. These indicate that *Slc39a8* deficiency in our model does not affect these measures of negative affective behavior at baseline. Previous investigation of similar phenotypes in neuronal-specific *Slc39a8* knockout models reported significant sex-specific effects for outer and intermediate zone durations (Choi et al., 2024), which we did not find in our study and could be due to different compensatory mechanisms that arise from the lack of this gene (Chen et al., 2018) and how those differ across tissues and models. Moreover, neuronal specific manipulations of this gene would account only for the investigation of its neuronal effects, masking its important blood brain barrier role (Liu et al., 2023).

The genetic influence on ethanol metabolism and its subsequent impacts on consumption and susceptibility to the development of AUDs have also been extensively studied and reviewed (Lehner et al., 2024; Edenberg and McClintick, 2019, Quertemont, 2004).

Furthermore, ethanol metabolites like acetaldehyde have been shown to be involved in reward pathway signaling, and thus can influence ethanol preference and consumption (Foddai et al., 2003; Melis et al., 2007). To assess whether metabolic clearance played a role in the differences we found in ethanol consumption, we tested BEC in our mice after 2 g/kg ethanol injections at different timepoints. No significant effect of genotype on BEC over time was found, suggesting that the changes in ethanol intake are not due to genotypic differences in metabolism. It is possible that measures of BEC following i.p. injections might not promote the same pharmacokinetic processes as voluntary oral intake, as ethanol absorption rates and bioavailability vary depending on delivery method (Pastino and Conolly, 2000). In addition, *Slc39a8* has been shown to have a role in digestive tissues (Li et al., 2016; Melia et al., 2019), making it possible that any genotype-dependent differences in absorption would not be seen in our current experimental design. Future studies could address this possibility by testing ethanol metabolism following oral gavage.

The A391T reduced-function variant has also been associated with increased body weight and BMI in human GWAS (Fernández-Rhodes et al., 2018; Haller et al., 2018; Locke et al., 2015). In our study, body weight data during experimental days revealed a significant genotype effect on body weight, with our HET mice showing higher weights than controls, potentially consistent with the direction of effect from human GWAS. Interestingly, several previous studies in different animal models of *Slc39a8-*deficiency have not seen a genotype effect on weight (Banna et al., 2022; Lin et al., 2017; Verouti et al., 2022), and one study found that their mutant SLC39A8-p.393T knock-in mice had lower weights than their controls during P14 and P30 stages (Li et al., 2022). Similar to our drinking data, these findings suggest that there may be subtle difference between different models of *Slc39a8-* deficiency, and care should be taken in choosing an appropriate strain that best models the phenotypes seen with human variants of *SLC39A8*.

In our study, we analyzed Zn^2+^ concentrations in blood, brain, and liver in naïve animals and after drinking in Experiment 1. We found trending levels for difference in Zn^2+^ levels in the brains of our naïve female mice, with HET animals exhibiting increased Zn^2+^ levels compared to that of WT. No differences were found in liver and blood samples. For the post-drinking samples, we found that HET animals showed significantly lower levels of Zn^2+^ in the liver. We also found a significant genotype effect for our blood samples, though we had a limited sample size and were unable to conduct the analysis with sex as a factor. A limitation in this experiment is that we were only able to analyze a small number of samples for some of the organs and our approach could not differentiate extracellular from intracellular Zn^2+^ pools and organ regional differences, which would potentially show greater genotypic differences. To address possible brain regional differences, we performed LA-ICP-MS in sagittal slices of WT and HET mice. Our results revealed higher concentrations of Zn^2+^ and Fe^2+^ in the ventral striatum of HET animals and trends towards genotype effects on prefrontal cortex and hypothalamus levels of Zn^2+^. Previous investigation of blood and brain Zn^2+^ and Fe^2+^ levels showed decreased Zn^2+^ levels in the brain cortex of A391T knock-in mice and increased blood levels, whereas no effect was seen in Fe^2+^ levels in either tissue (Li et al., 2022). Tseng et al. showed reduced Zn^2+^ transport in cell models with the SLC39A8-p.391T variant (which corresponds with the rs13107325 and A391T variants). Lin et al. found no effects of inducible global deletion of *Slc39a8* in Zn^2+^ and Fe^2+^ levels in several tissues, whereas Zhang et al. highlight the role of this gene in Fe^2+^ recycling and distribution. Previous work using an inducible model did not find brain differences in either Zn^2+^ and Fe^2+^ levels (Lin et al. 2017). A common limitation across studies is the lack of regional specificity when analyzing brain levels of these metals. This emphasizes the strength of methods such as LA-ICP-MS, which allows for the measurement of element distributions across much smaller brain regions and may reveal subtle genotypic differences that are obscured when analyzing whole brain.

In the context of our study and findings, it has been demonstrated that Zn^2+^ concentrations modulate the effects of ethanol on glycine receptors (McCracken et al., 2013), which in turn regulate ethanol-dependent dopamine release and sensitivity in the nucleus accumbens (Loftén et al., 2023; San Martin et al., 2021). These indicate that increases in brain Zn^2+^ availability could increase sensitivity to ethanol due to glycine receptor potentiation and, as a consequence, limit ethanol intake. Additionally, *Slc39a8* deficiency promotes differential surface expression of NMDA receptor subunits (Tseng et al., 2021), and these have repeatedly been implicated in different ethanol-related effects in different brain regions (Suvarna et al., 2005; Lovinger, 1995; Popp et al., 1999; Jury et al., 2019; Zamudio et al., 2022). To our knowledge, this is the first study to investigate striatal metal concentrations in models of *Slc39a8* deficiency. The dopamine transporter (DAT), which is essential in striatal functioning, has been shown to have an endogenous Zn^2+^ binding site (Norregaard et al., 1998). Differential Zn^2+^ content in the striatum has been shown to have several regulatory effects on dopamine reuptake, functioning both in increasing and decreasing DAT function, depending on membrane potential and on the presence of other ions (Li et al., 2015; Pifl et al., 2009) and it also has behavioral outcomes upon cocaine exposure, such as potentiation of cocaine locomotor sensitization and reward seeking (Gomez et al., 2021). Zn^2+^ and *Slc39a8* are also both involved in immune regulation, which could potentially contribute to our observed differences in ethanol consumption as increasing evidence supports the importance of the neuroimmune system in drug-associated behaviors. *Slc39a8* has been shown to act as a negative feedback regulator in the NF-κB pathway. At the onset of inflammation, *Slc39a8* is upregulated following NF-κB activation and subsequently downregulates the NF-κB cascade through inhibition of IKKβ by promoting Zn^2+^ influx into the cytosol (Liu et al., 2013). Activation of NF-κB in the ventral striatum was found to be necessary for neuro-immune alterations that facilitate cocaine self-administration in stress-induced models (Mongi-Bragato et al., 2025), and antagonism of the same pathway was shown to result in decreased ethanol preference (Montesinos, Gil, and Guerri, 2017). These demonstrate general examples of how different mechanisms influenced by Zn^2+^ and *Slc39a8* could impact consumption behavior in our *Slc39a8* deficient mice. Future mechanistic studies will be needed to test these possibilities directly. Moreover, given the high importance of trace metal homeostasis for proper biological functioning, compensatory mechanisms to maintain Zn^2+^ homeostasis in the HET mice should also be explored.

Different variants of *Slc39a8* have been repeatedly associated with AUD and other ethanol-related traits in GWAS, highlighting the importance of further investigation to better understand the role of this gene in these phenotypes. Specifically, the missense A391T variant has been repeatedly associated with lower levels of drinking (Liu et al., 2019), which is consistent with the direction of effect of heterozygous knockout on ethanol consumption see here in a mouse model. Our characterization of other phenotypes in this model provide evidence that differences in ethanol consumption are not due to metabolic changes or altered rewarding solution consumption. Finally, our initial investigation of brain and other tissue levels of Zn^2+^ suggests a possible mechanism underlying the lower ethanol intake seen with reduced *Slc39a8* function. These studies may pave the way for future examination of reduced function *Slc39a8* variants as protective factors against AUD, and may inform novel insights into the underlying neurobiology of AUD and possible therapeutic strategies.

## Data availability

The data collected and analyzed in this study are available on reasonable request from the corresponding author.

## Acknowledgements

This work was supported by a grant from the NIH-NIAAA to AMB-L (R00AA027835). LA-ICP-MS experiments were supported by the University of New Mexico Center for Metals in Biology & Medicine through NIH-NIGMS grant P20GM130422.

## Author contributions

ALS-B: conceptualization, data collection and analysis, and writing the original draft of the manuscript. D-JGP, RS, and CGP: conceptualization, data collection, and review and editing of the manuscript. AMB-L: conceptualization, supervision, funding acquisition, and review and editing of the manuscript.

## Competing interests

The authors declare no conflicts of interest.

## Notes

### Competing Interest Statement

The authors have declared no competing interest.

