## Supplement Figures for "Functional validation and characterization of *Slc39a8* genotype effects on ethanol consumption and related phenotypes"

Day 1

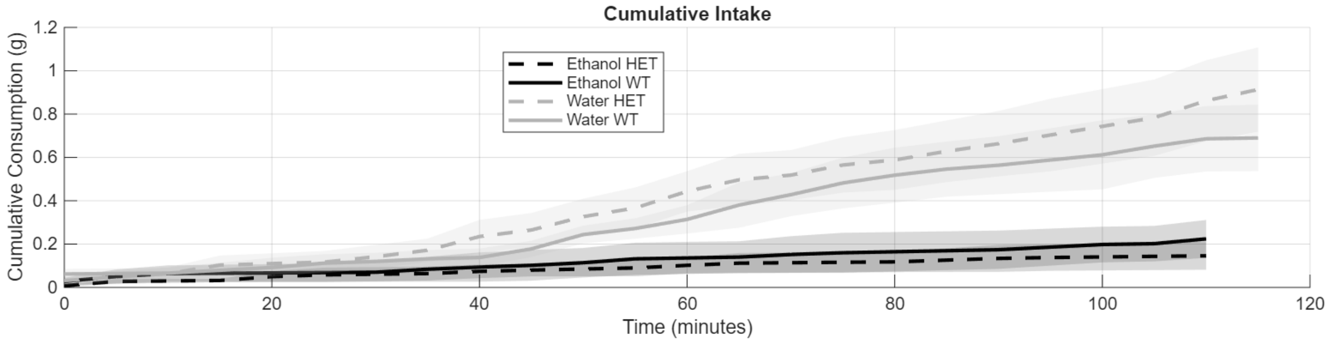


Day 2

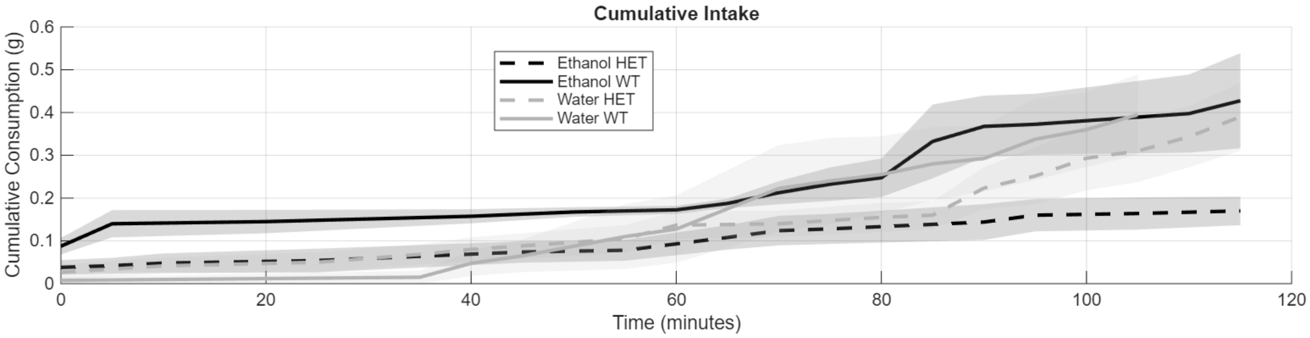


Day 3

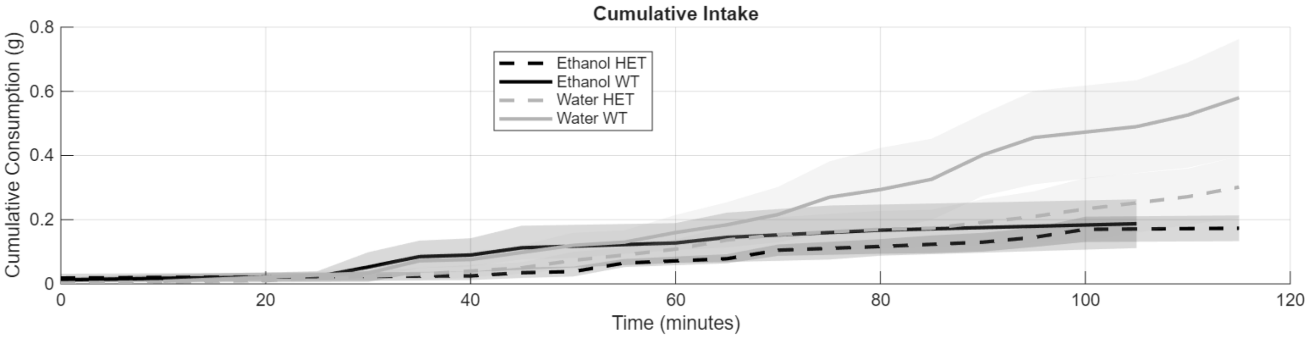


Day 4 0-2h

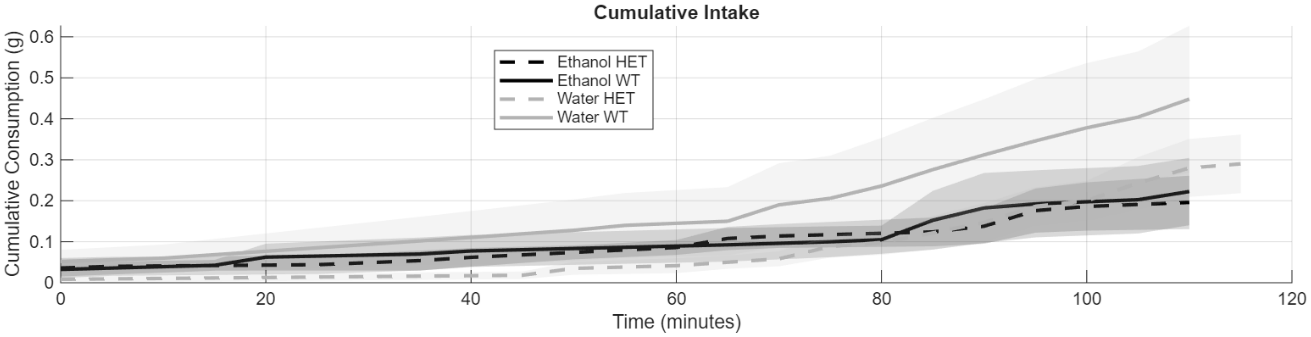

Day 4 2-4h

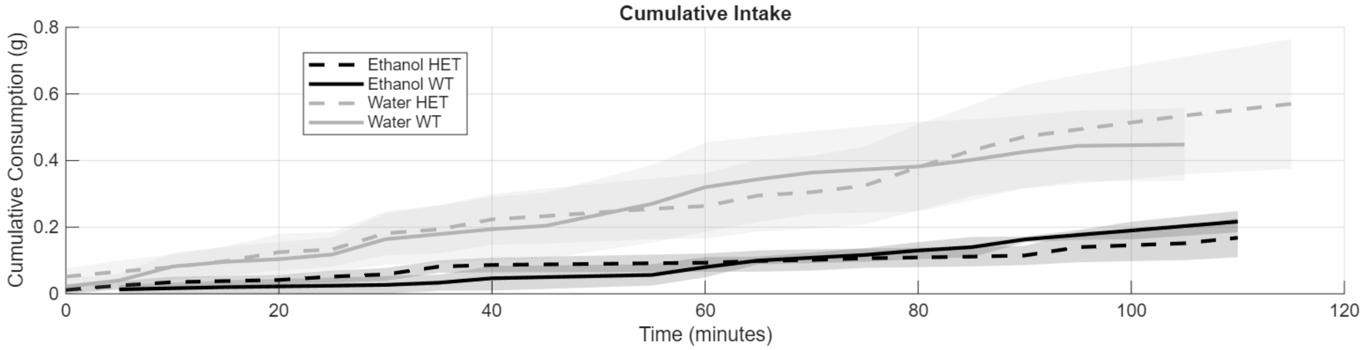


Day 4 24h

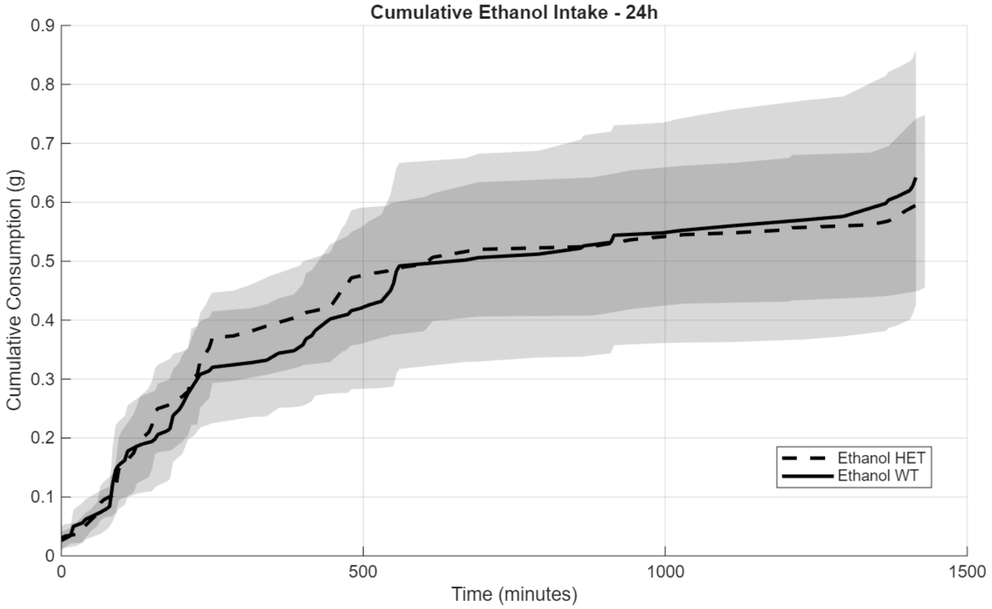


Supplementary Figure S1: Cumulative fluid consumption during two-bottle choice DID days. N= 5-6/genotype.

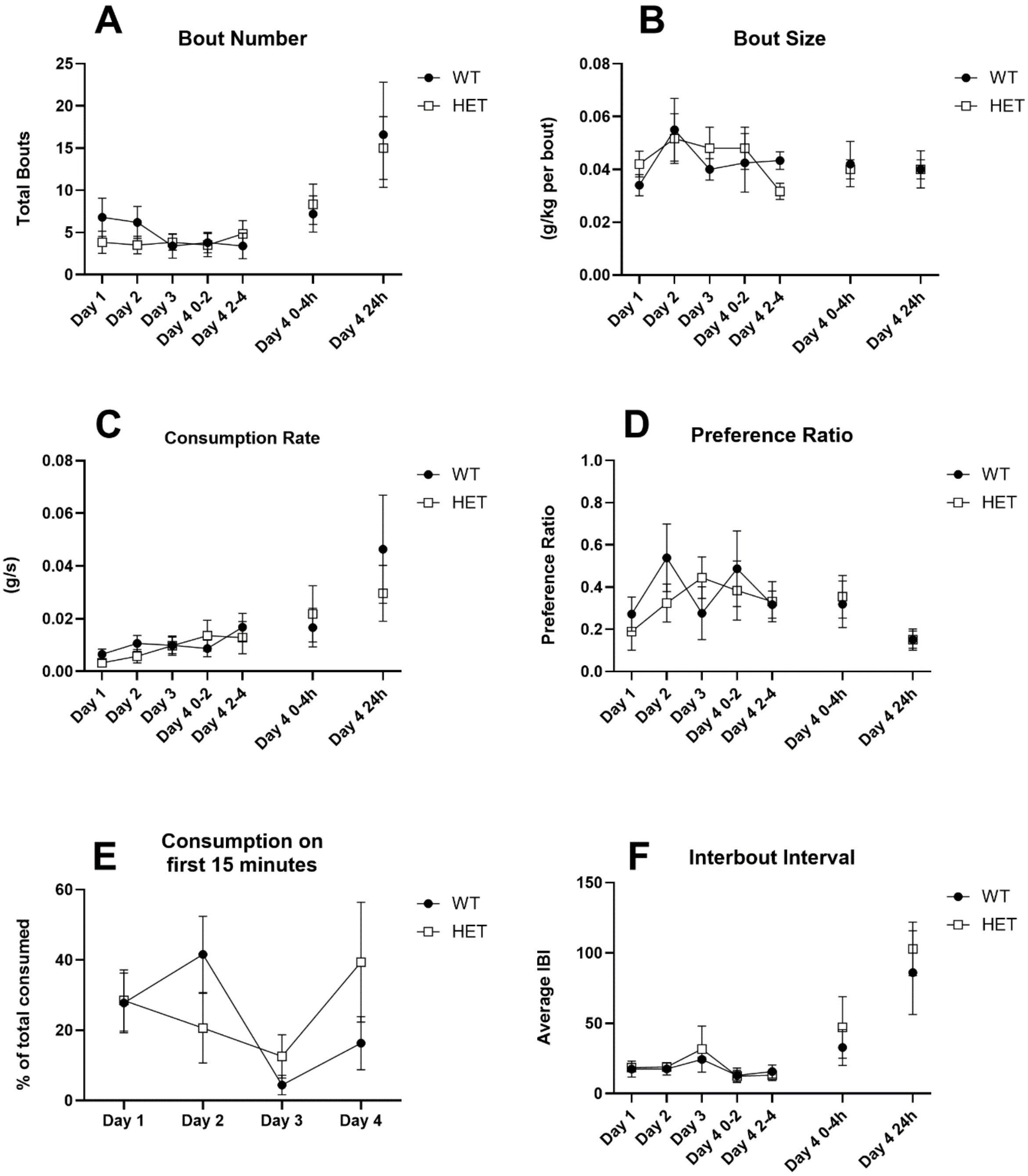


Supplementary Figure S2: Two-bottle choice DID in the BioDAQ. Panels show bout number (A) and size (B), ethanol consumption rate (C), preference ratio (D), percent consumed during first 15 minutes of each session (E), and interbout interval (F). Data are shown as mean ± SEM. N= 5-6/genotype.

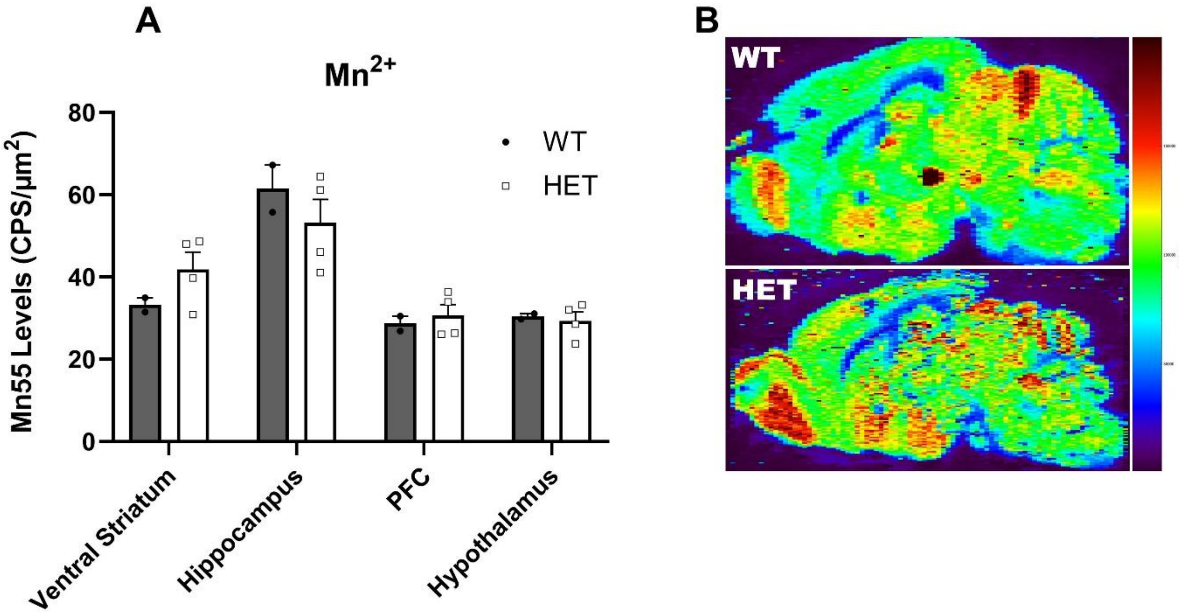


Supplementary Figure S3: Semiquantitative levels of $Mn^{2+}$ in different brain regions of naïve *Slc39a8* HET mice and their WT littermates (A) and representative elemental distribution map of $Mn^{2+}$ levels from the LA-ICP-MS analysis (B). Data are shown as mean ± SEM. N= 2-4 per genotype.

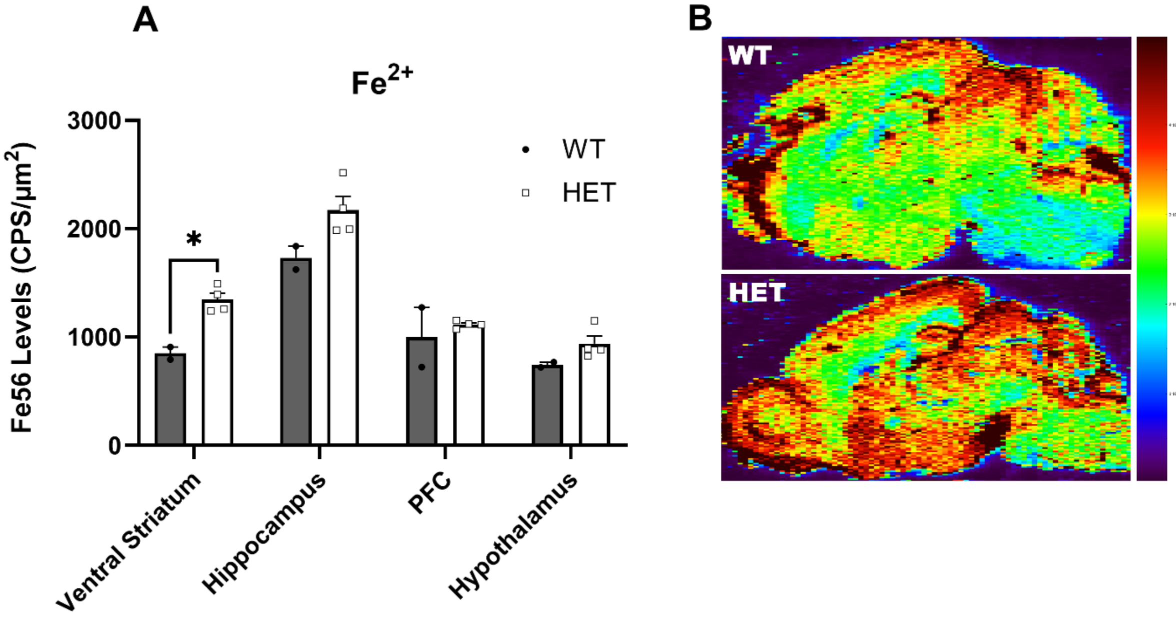


Supplementary Figure S4: Semiquantitative levels of $Fe^{2+}$ in different brain regions of naïve *Slc39a8* HET mice and their WT littermates (A) and representative elemental distribution map of $Fe^{2+}$ levels from the LA-ICP-MS analysis (B). Data are shown as mean ± SEM. * Indicates $p<0.05$. N= 2-4 per genotype.
